# Structural features recapitulate collective dynamics of inhibitory networks

**DOI:** 10.1101/2019.12.17.879726

**Authors:** Shivik Garg, Divye Singh

**Affiliations:** Computational Neurobiology Laboratory - I, Indian Institute of Science Education and Research Pune, Dr Homi Bhabha Road, Pune - 411008; Centre for Modeling and Simulation, Savitribai Phule Pune University, Ganeshkhind Road, Ganeshkhind, Pune - 411007

**Keywords:** Clustering, Inhibition, Olfaction, Spatiotemporal patterns

## Abstract

Inhibitory interneurons are ubiquitous throughout the central nervous system (CNS) and play an important role in organizing the excitatory neuronal populations into spatiotemporal patterns. These spatiotemporal patterns are believed to play a vital role in encoding sensory information. The olfactory system is a wellknown example where odor information is encoded in temporally evolving activity of the principal neurons and inhibitory interneurons play an important role in generating these patterns. In this work we study how inhibitory interactions generate such patterns in the context of odor encoding by simulating random biophysical models of mitral cells. Using the Newman community clustering algorithm we identify synchronously firing groups of neurons that switch in their activity. Our study presents a new method of inferring the dynamics of inhibitory networks from their structure.

## 1 Introduction

Spatiotemporal patterning of neural activity is known to play an important role in many functions like spatial navigation (Wilson and McNaughton (1993)), memory consolidation (Ji and Wilson (2007)),executing motor movements (Marder and Calabrese (1996)) and odor encoding (Laurent (2002); Laurent et al. (1996)). These temporally evolving patterns are dependent upon the structural distribution and different time scales of inhibition. In the hippocampus, the interneuronal network can organize the pyramidal cells into synchronously firing temporal sequences (Wang and Buzsaki (1996)). Central pattern generators (CPGs) which give rise to rhythmic activity consist of reciprocally inhibiting neurons (Marder and Calabrese (1996)). The olfactory circuit is a well-known example of a system where the temporally evolving activity of the principal neurons encodes odor information (Laurent (2002)). The olfactory circuit consists of a large population of inhibitory interneurons (granule cells) which far exceed in number the excitatory principal neuron population (mitral cells). These interneurons make reciprocal dendrodendritic connections with the mitral cells and help in shaping their dynamics. The synchronization of the principal neurons (mitral cells) occurs via the inhibitory synaptic inputs from the granule cells (Schoppa (2006)). In honeybees, blocking GABAergic inhibition in the antennal lobe (equivalent to the mammalian OB) by picrotoxin leads to desynchronization of the principal neurons and they fail to discriminate between structurally similar odor molecules (Stopfer et al. (1997)). Increasing inhibition on to the mitral cells leads to faster discrimination between similar odor pairs (Abraham et al. (2010)). Computational modeling of the locust antennal lobe has shown that the local interneuron (LN) subnetwork entrains the PN population giving rise to spatiotemporal patterning (Bazhenov et al. (2001)). In this article, we examined how the olfactory bulb network give rise to such evolving cell assemblies (spatiotemporal patterning) of mitral cells. We addressed this question using randomly connected (via lateral inhibition), biophysi cally spiking mitral cell models.Previous work has shown that randomly connected inhibitory networks generate sequentially firing cell assemblies accounting for the observed striatal network dynamics (Ponzi and Wickens (2010)) Identifying groups of synchronously firing neurons in inhibitory networks is a challenging problem. Assisi et. al studied inhibitory network dynamics by relating it to a structural property of the network, called coloring (Assisi et al. (2011)). Graph coloring is useful for smaller and more regular networks, however, it cannot be used for random graphs as there are various possible minimal colorings and determining them in a reasonable time is a difficult problem. In this study, we overcome these limitations by using the Newman community clustering algorithm to explore the relationship between the structure of inhibitory networks and their dynamics in the context of odor encoding.

## 2 Methods

### 2.1 Mitral cell and synapse model

Mitral cell and synapse model: In this study, we used realistic conductance based, single-compartment models of mitral cells. The model consists of spiking sodium and potassium currents along with a transient potassium current which causes a delay in the onset of the spike. The model also has channels for slow potassium and persistent sodium current which, together, give rise to subthreshold oscillations. The membrane potential for each mitral cell is given by the following equation:

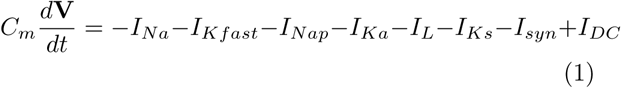

The parameters for the mitral cell model were adapted as previously described in Bathellier et al. (2006). Below we list the parameters that have been modified from previous work:

IKa current:

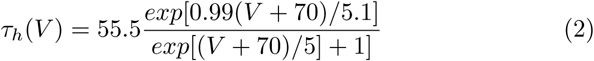

IKs current:

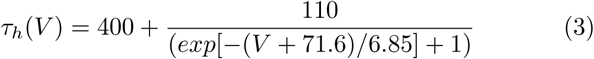

Synaptic conductance is modeled as a difference of two exponential functions of time t

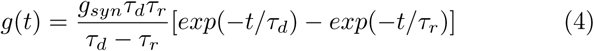

where *τ*_*d*_ and *τ*_*r*_ are the rise time and fall time of the synapse respectively. *g*_*syn*_ was taken to be 5.2381, because of this the maximal value of g(t) is 1 with respect to time. The synaptic current is given by :

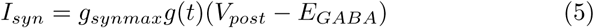

where *g*_*synmax*_ is the maximal synaptic conductance. The inhibitory synaptic current is activated by presy-naptic spikes in the mitral cell. The reversal potential for GABAergic synapse is -70.0 mV. The rise time is 0.2 ms and the decay time is 20 ms. Each synapse gets activated with a latency of 0.75 ms. For results presented in Fig. 2 *g*_*synmax*_ = 6 *mS/cm*^2^ and for Fig. 3 *g*_*synmax*_ is normalized based on the degree of connectivity of the neuron.

**Fig. 1.**
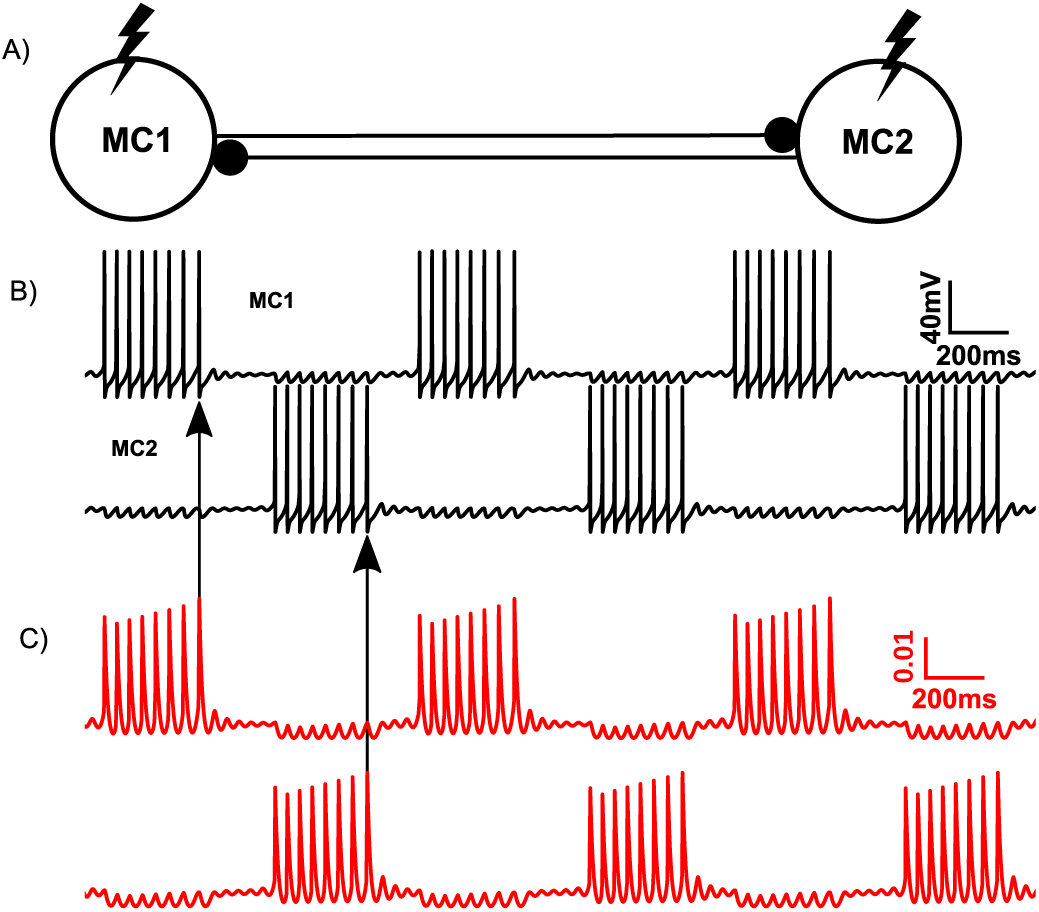
Dynamics of two reciprocally inhibiting mitral cells. **A)** Schematic of two mitral cells reciprocally inhibiting each other. **B)** Reciprocally inhibiting mitral cells when driven by depolarizing input show alternation in their activity. **C)** The traces represent the open fraction of slow potassium current. Build-up of the slow potassium current causes the cell to stop firing leading to a switch in their firing activity.

**Fig. 2.**
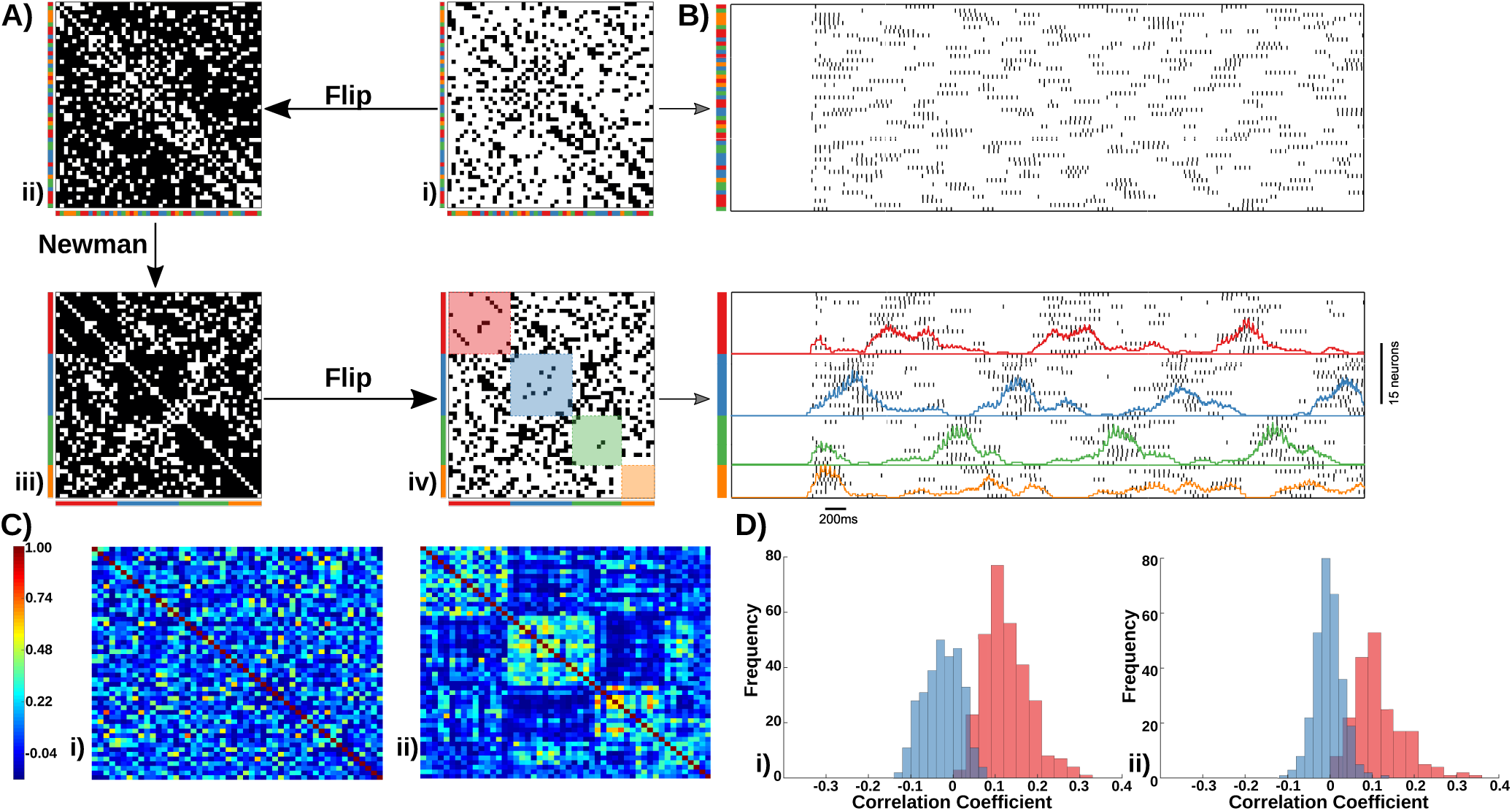
Clustering of random inhibitory networks. **(A)** Schematic showing how groups of neurons which are maximally disconnected with each other are found for a particular random network. The square boxes represent connectivity matrices (black represents connection whereas white indicates no connection). **(B)** Raster plot showing the activity of the neurons of the original connectivity matrix (top panel) and after rearranging the neurons according to the groups obtained after clustering (bottom panel). The colored traces show the averaged activity of each group. **(C)** Correlation matrices of activity of 50 neurons. Panel i) shows the correlation coefficient for a randomly clustered network and panel **(ii)** shows the correlation coefficient for the network permuted based on the Newman algorithm. In panel **(ii)** there is the presence of a group like structure which is absent in panel **(i). (D) i)** shows the distribution of within-group (red) and across group correlations (blue) for different noise trials of a particular random network. **ii)** shows the same distribution for different random networks.

**Fig. 3.**
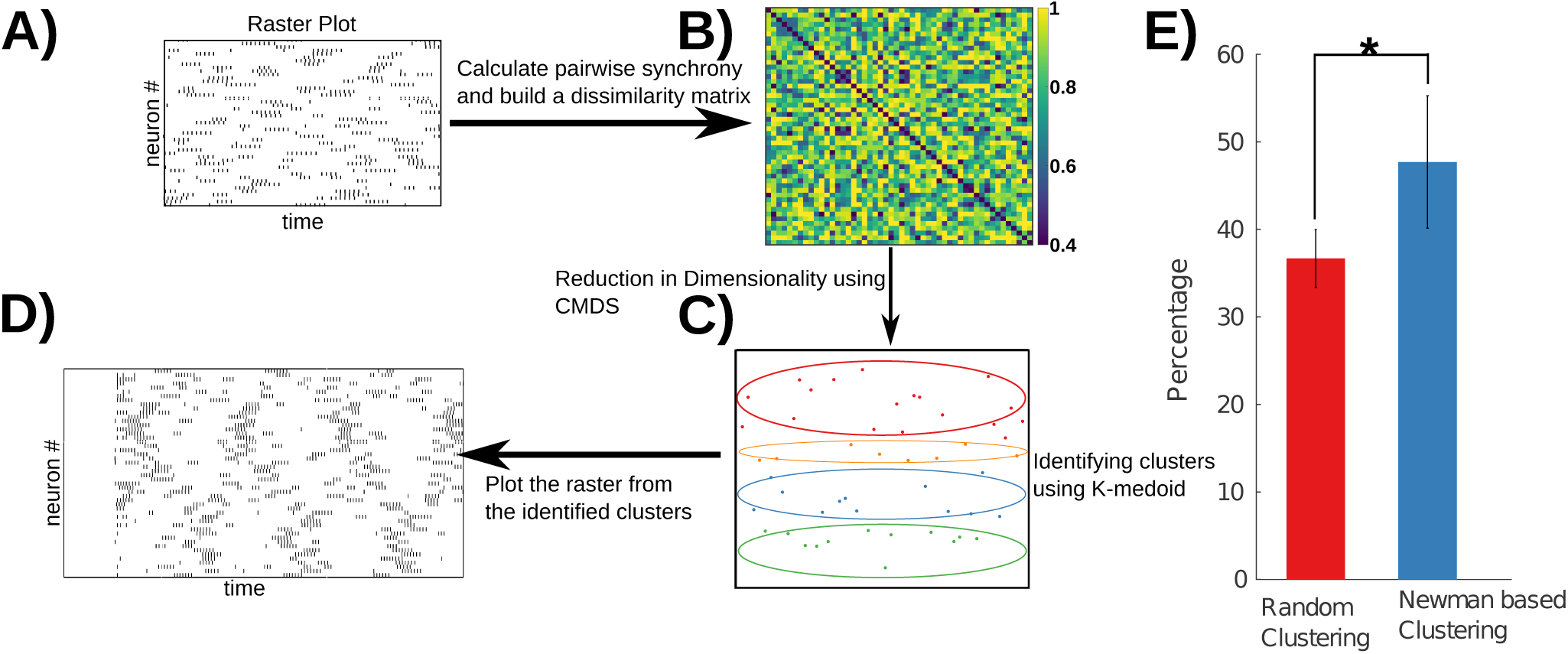
Clustering based on the dynamics. Overall schematic for clustering network based on the dynamics. Spike times of individual neurons are recorded **(A)** which are then used to calculate pairwise spike synchrony. The synchrony measure is then used to construct a dissimilarity matrix **(B)**. This dissimilarity matrix is further used to identify clusters using the k-medoid algorithm. The dimensionality of this system is further reduced to a single dimension using Classical Multi-Dimensional Scaling (CMDS) to visualize the clusters identified **(C)**. The groups obtained through this method were used to replot the dynamics (shown as a raster plot) **(D). (E)** Bar plot showing the maximal overlap between randomly clustered and dynamics based clustering and Newman based clustering and dynamics based clustering. Overlap of the neurons in the clusters obtained from the dynamics based clustering with the Newman based clustering is significantly better when compared to different random clusterings (two-tailed Student’s t-test p *<* 0.01).

### 2.2 Network Structure

We constructed random networks using the Watts and Strogatz model. The random networks used for simulations consisted of 50 neurons and the results presented are for 50 randomly generated networks. The number of nearest neighbour connections (K) was set to 6 and the rewiring probability (p), which sets the randomness of the network, was 0.8 (Watts and Strogatz (1998)).

### 2.3 Newman Community Clustering

The Newman community clustering algorithm detects communities based on edge densities such that the maximum number of edges lie within a group and minimum number of edges lie across groups. We implemented the Newman community clustering algorithm from the Brain Connectivity Toolbox in MATLAB (Rubinov and Sporns (2010)).

### 2.4 Classical Multidimensional scaling (CMDS) and K-medoid clustering

CMDS uses eigenvalue decomposition for dimensionality reduction. K-medoid is a partitioning algorithm used to cluster data sets into different groups. In this method the aim is to group data into clusters such that their distance to the medoid of the cluster is minimized. Both methods were implemented using R.

### 2.5 Spike Synchrony

To calculate spike synchrony we used a measure called SPIKE synchronization. This was implemented using a python library known as ‘pyspike’ (Mulansky and Kreuz (2016)).

### 2.6 Correlation coefficient

The pairwise correlation coefficient was calculated for the firing rate obtained over a 50 ms window. Since the number of clusters obtained for different networks varied we calculated correlations only for the highest three groups.

### 2.7 Simulation procedure

All network simulations were carried out using a fourth-order Runge Kutta integration method. The time step used for simulation was 27 *µs*. The input onset was set at 1000 ms so that the cells settle down to steady-state. The input (DC of 0.6 *µ*A/cm^2^) was provided for 6000 ms and started to decay at 7000 ms. The entire simulation was done for 8000 ms. Simulations were performed using an in house developed open-source C++ library called ‘in-silico’ (http://insilico-lib.github.io/insilico/). The in house ‘in-silico’ utilizes a boost library (odeint) for integrating systems of coupled differential equations. MATLAB was used to analyze the data.

## 3 Results

### 3.1 Mutually inhibitory motif of two mitral cells

The olfactory bulb circuitry consists of mitral cells coupled to each other via the inhibitory interneurons (granule cells) (Shepherd and Greer (2004)). Mitral cells (the principal neurons in the olfactory circuit) mediate lateral inhibition onto each other through these inhibitory interneurons. To examine the effects of inhibition, we first simulated a simple network motif consisting of two mitral cells. The granule cells were modeled implicitly by implementing a direct GABAergic synapse (lateral inhibition) between the two mitral cells. The two mitral cells, when driven by an identical depolarizing current, showed switching in their spiking activity (Fig. 1). This transition in the activity occurred because of the slow potassium current. The slow potassium current is a depolarization activated hyperpolarizing current that builds slowly with each successive spike, eventually causing the mitral cell (MC1) to cease firing. This releases the other mitral cell (MC2) from inhibition and it starts firing and the whole cycle of events repeats.

### 3.2 Dynamics of random inhibitory networks

Since mitral cells coupled through reciprocal inhibition showed antagonistic interactions, we attempted a novel method to study the dynamics of larger networks. In these networks, we looked for groups of neurons such that neurons belonging to a particular group would be maximally disconnected from each other. We posited that such a group of neurons would fire synchronously whereas neurons associated with different groups would fire at different times. We employed the Newman algorithm (Newman (2006)) to study this relationship between the structure of the network and its dynamics. We constructed random networks of mitral cells inhibiting each other (see METHODS). To find the group of neurons that are maximally disconnected from each other we flipped the connectivity matrix by converting 0’s to 1’s and the 1’s to 0’s (no self connections present)(Fig. 2A (i)) and passed it through the Newman algorithm (Fig. 2A (ii)). The algorithm is then used to detect clusters of neurons in the inverted connectivity matrix. Using these communities, we rearranged the inverted connectivity matrix such that neurons belonging to a particular cluster are grouped together (Fig. 2A (iii)). We then flipped this matrix again to arrive at a new permutation of the original connectivity matrix.

This operation revealed a block-like structure which showed that most of the connections were present on the off-diagonal (Fig. 2A (iv)). In the example shown, the reordered connectivity matrix reveals the presence of 4 groups (color-coded as red, blue, green and orange). The dynamics of the original connectivity matrix (plotted as a raster) did not show any temporal patterning. However, when we plotted the dynamics of the reordered connectivity matrix we find that a temporal patterning exists (cell assemblies that switch in their activity) (Fig. 2B). Neurons associated with each color fired synchronously and were anti-correlated with the other groups. The correlations in the firing rates showed the presence of a group like structure when compared with a random clustering of the network, which did not show any such clustering (Fig. 2C). These correlations persisted for simulations of a particular random network across different noise trials (Fig. 2D (i)). Similar results were obtained for simulations of several randomly generated networks (Fig. 2D (ii)).

### 3.3 Dynamics based clustering of random networks

We then asked whether we could cluster the neurons using their firing activity and if this would correlate with the clustering obtained from the Newman algorithm. We recorded the spike times of all the neurons (Fig. 3A) and calculated pairwise synchrony using SPIKE synchronization, which was then used to create a dissimilarity matrix (by subtracting them from 1) (Fig. 3B). Clusters were then detected using k-medoid clustering method (Fig. 3C). Replotting the activity (as a raster) after clustering (Fig. 3D) revealed the underlying antagonistic interactions between groups, showing the presence of a spatiotemporal pattern. We further calculated the overlap between the groups obtained from Newman clustering and dynamics based clustering. Since the identities of the groups found through the dynamics based clustering are not known, we calculated the best overlap between Newman based clustering and structure-based clustering. We calculated the overlap for all possible combinations and the highest overlap value was chosen for a particular random network. Fig. 3E shows that dynamics based clustering had a greater overlap with the Newman based clustering when compared to a random clustering.

## 4 Discussion

In this study, we used a new approach to study the relationship between the structure of inhibitory networks and their resulting dynamics. Using a community clustering algorithm (Newman (2006)) we arrived at a partitioning of the network, such that neurons belonging to a group are maximally disconnected from each other but connected to neurons belonging to other groups. Using this definition we were able to extract correlations between the network dynamics and its structure. Further we identified groups of neurons firing synchronously based purely on their dynamics. We found that the resulting clusters overlapped significantly with the groups obtained based on the network structure thus revealing that our approach of predicting the dynamics based on structure was reliable. Our approach allowed us to generate a lower-dimensional representation of a complex high-dimensional system. This approach also allows for devising new ways of inferring the connectivity of the network from the dynamics.

Studies of the insect antennal lobe (AL) and the olfactory bulb (OB) have shown that circuit interactions within the AL/OB generate the temporal structure of the principal neuron activity and inhibition plays an important role in generating these patterns (MacLeod and Laurent (1996)). Our simulations support the role of lateral inhibition in generating these patterns among the mitral cell population (Friedrich and Laurent (2001)). One caveat of our work is that we have used a reduced olfactory bulb circuitry (a purely inhibitory network of mitral cells), whereas they are actually coupled by granule cells.

Sequential firing of cell assemblies have been proposed as a mechanism of information encoding by the nervous system for downstream targets. Theoretical work has shown that the origin of such dynamics occurs because of the winnerless competition (WLC)(Rabinovich et al. (2001)). Spatiotemporal patterning in our case also seems to occur because of this mechanism. Our work complements previous work where lateral inhibition in random networks predictably leads to the formation of cell assemblies (Ponzi and Wickens (2010)). Inhibitory interactions amongst neurons are known to play an important role across different anatomical regions in the CNS (Marder and Calabrese (1996); Buzsaki and Chrobak (1995)). Inhibition can be used to synchronize excitatory neuronal populations and organize them into events of sequential activity. These sequences could potentially be used by the nervous system to encode information in a more reliable and condensed form.

## Acknowledgements

We would like to thank Dr. Collins Assisi for fruitful discussions.

